# *Mx1*-labeled pulp progenitor cells are main contributors to postnatal odontoblasts and pulp cells in murine molars

**DOI:** 10.1101/2024.03.21.586156

**Authors:** Dongwook Yang, YoungJae Jeong, Laura Ortinau, Jea Solidum, Dongsu Park

## Abstract

Regeneration of dentin and odontoblasts from dental pulp stem cells (DPSCs) is essential for permanent tooth maintenance. However, the *in vivo* identity and role of endogenous DPSCs in reparative dentinogenesis are elusive. Here, using pulp single-cell analysis before and after molar eruption, we revealed that endogenous DPSCs are enriched in *Cxcl12-*GFP^+^ coronal papilla-like cells with *Mx1-*Cre labeling. These *Mx1*^+^*Cxcl12-*GFP^+^ cells are long-term repopulating cells that contribute to the majority of pulp cells and new odontoblasts after eruption. Upon molar injury, *Mx1*^+^ DPSCs localize into the injury site and differentiate into new odontoblasts, forming *scleraxis*-GFP^+^ and *osteocalcin*-GFP^+^ dentinal tubules and reparative dentin. Single-cell and FACS analysis showed that *Mx1*^+^*Cxcl12-*GFP^+^ DPSCs are the most primitive cells with stem cell marker expression and odontoblast differentiation. Taken together, our findings demonstrate that *Mx1* labels postnatal DSPCs, which are the main source of pulp cells and new odontoblasts with reparative dentinogenesis *in vivo*.

## INTRODUCTION

All human teeth are permanent and, therefore, the natural regeneration and repair of tooth cells and mineral layers are essential for the long-term maintenance of dental tissues. In particular, dental pulp cells and differentiated dentin-producing odontoblasts play an important role in supporting tooth structure, absorbing the mechanical pressure from mastication and sensing thermal, osmotic, and tactile stimuli^1^. When a tooth is exposed to wear, malignant caries, and fractures, the pulp and odontoblasts contribute to the development of a protective barrier, dentin regeneration, and immune responses. By contrast, severe pulpal damage or endodontic treatment can cause loss of sensation, increased fragility, and tooth loss. Therefore, proper maintenance and regulation of pulp progenitor cells and their differentiation to new odontoblasts are important for permanent tooth health and maintenance. Previously, external stimuli such as uniaxial tensional/compressional stress have been reported to induce progenitor proliferation and odontogenic differentiation^2–5^. Furthermore, a recent odontoblast depletion study showed that new odontoblast-like cells can be regenerated from the cell-rich zone of the pulp, suggesting the presence of progenitor cells in adult molar pulp^6^. However, due to the heterogeneity of pulp cells and the limited availability of animal models to label endogenous pulp progenitor cells, it is largely unknown which cells are dental pulp stem cells (DPSCs) and how they contribute to endogenous odontoblast regeneration.

During tooth development, *Wnt1*-Cre^+^ neural crest cells were known to be a source of primitive pulp (papilla) cells and odontoblasts for primary and secondary dentin formation^7–9^. However, in permanent teeth, pulp cells near odontoblasts were reported to proliferate and differentiate into odontoblast-like cells in response to odontoblast depletion^6,8,9^. Other studies showed that *Axin2*^+^ cells or *aSMA*^+^ perivascular cells in the dental pulp expanded through proliferation and gave rise to odontoblast-like cells, suggesting that adult DPSCs reside in the tooth pulp^10,11^. By contrast, lineage tracing of *Sox10*-CreER^+^ and *PLP*-CreER^+^ cells with a Schwann cell origin showed that glial cells generate pulp progenitor cells and contribute to odontoblasts in adult growing incisors^12^. Furthermore, recent studies revealed that *Gli1*^+^ cells and PTHrP^+^ cells in the developmental tooth apical region (PDLSCs) give rise to PDL, alveolar bone, and a subset of pulp cells, but do not contribute to postnatal pulp cells and odontoblasts, indicating the presence of multiple pulp progenitor cells^13–15^. In fact, the labeled cells in many of these studies were heterogeneous and their tooth injury models caused pulp exposure and the destruction of adjacent dentin structures. Therefore, it is still not clear whether existing pre-odontoblasts or new odontoblasts from stem/progenitor cells replace injured odontoblasts during the regeneration process. Moreover, it is essentially unknown which of the pulp cells are long-term DPSCs and whether they contribute to the recycling of adult pulp cells and odontoblasts in permanent teeth.

In this study, we sought to define the molecular characteristics and function of endogenous DPSCs in permanent molar teeth and found that the *Mx1* and *Cxcl12-*GFP combination can be induced to selectively label endogenous DPSCs. We further sought to define how *Mx1*^+^ progenitor cells respond to external stimuli and contribute to the replacement of odontoblasts and found that *Mx1*^+^ progenitor cells are major contributors to postnatal pulp cells and supply new odontoblasts with reparative dentinogenesis under homeostatic and injury conditions.

## RESULTS

### Pulp cells in post-erupted molar tooth acquire coronal papilla-like characteristics and highly express Cxcl12

Dental pulp cells are known to have a neural crest cell (NCC) origin^16,17^. However, adult pulp cells and DPSCs have a unique environment with less defined endogenous function and identity. In particular, how the population and molecular characteristics of pulp cells change before and after eruption of a molar tooth has not been studied.

To define molecular changes in pulp cells before and after tooth eruption, we performed single cell RNA sequencing of the mouse first molar at P25 (post-eruption stage), and normalized and compared the results to previously reported datasets from the pre-eruption (P3.5) and eruption (P7.5) stages of the mouse first molar tooth (Figure 1A). UMAP clustering analysis of integrated datasets showed that molar teeth comprised at least 10 distinct clusters (Figure 1B). Among these clusters, cluster 1 selectively expressed early coronal papilla markers (Enpp6, Fabp7, Fmod), cluster 2 expressed mid papilla markers (Nnat4, Rab3b, Vcan), and cluster 3 expressed apical papilla markers (Aox3, Tac1, Crabp1), annotating that cluster 1, 2, and 3 represent dental papilla cells that can develop into pulp cells and odontoblasts. An additional cell-type specific marker analysis revealed that cluster 4 expressed odontoblast markers (Ocn, Dmp) and cluster 5 and 6 expressed dental follicle cell markers, while epithelial and ameloblast markers were enriched in cluster 7 and 8 and endothelial and immune cell markers were highly expressed in cluster 10 and 11, respectively (Figures 1B and S1A). Next, we separately analyzed pre-eruption (P3.5), eruption (P7.5) and post-eruption (P25) datasets and examined pulp populational changes between pre- and post-eruption of the permanent tooth. Notably, a majority of pulp cells from the post-eruption stage (P25) showed a marked decrease of apical and middle papilla markers and switched to Enpp6^+^ coronal papilla (CP)-like cells (Figures 1C and S1A). In addition, these P25 CP-like subsets showed a significant increase in the expression of progenitor cell markers, CXCL12 and PDGFR, compared to P3.5 and P7.5 CP subsets (Figures 1C and 1D). However, as invariable controls, mature odontoblasts (cluster 4) and endothelial cells (cluster 10) did not change their numbers and marker expression throughout all time points (Figures 1C and S1B). Consistently, differential gene analysis and pseudo-time analysis of P25 pulp cells revealed that the CP-like cluster (cluster 1) at P25 highly expressed stem cell markers (Cxcl12, Enpp2, Ifitm1) and positive regulators of odontoblastic differentiation (i.e. Slc20a2, Cldn10, Bmp2, Timp3), but decreased the expression of early apical papilla markers Notum and Smpd3 (Figures 1D-1E and S1C)^13,18–28^. These results suggest that, unlike incisors, molar pulp progenitor cells undergo a unique population change after eruption and acquire the characteristics of coronal papilla cells with high CXCL12 expression.

**Figure 1.**
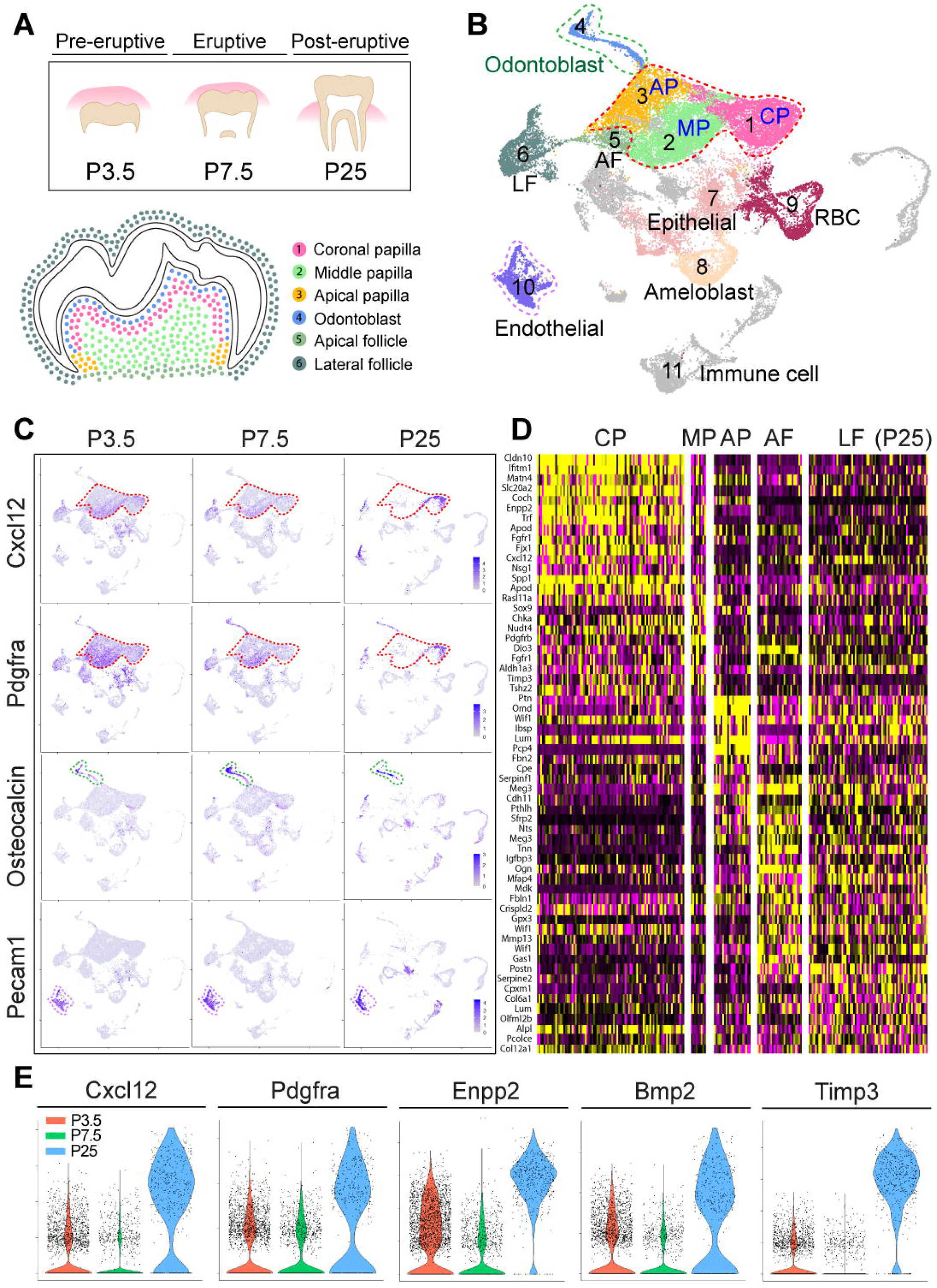
Single-cell analysis of pre- and post-erupted molars reveals unique clusters of postnatal pulp progenitor cells. (A) Schematic diagram of molar eruption and single-cell sequencing from post-erupted first molars (P25) compared with previously reported datasets from pre-eruption (P3.5) and eruption (P7.5) stages. The bottom figure is a schematic diagram identifying clusters present in the tooth germ in the 1st molar of the pre-eruptive stage. (B) Integrated UMAP visualization of 11 different color-coded clusters; Cluster 1: Coronal papilla (CP), Cluster 2: Middle papilla (MP), Cluster 3: Apical papilla (AP) (all papilla clusters with blue dots), Cluster 4: Odontoblast (green dots), Cluster 5: Apical follicle (AF), Cluster 6: Lateral follicle (LF), Cluster 7: Epithelial, Cluster 8: Ameloblast, Cluster 9: Red blood cell (RBC), Cluster 10: Endothelial (purple dots), Cluster 11: Immune cell. (C) UMAP-based transcriptional plots for Cxcl12, Pdgfra, Osteocalcin, and Pecam1 at P3.5, P7.5, and P25. Dashed lines indicate the coronal papilla (red), odontoblast (green), and endothelial (purple) clusters at each time point. (D) Heatmaps showing the top differentially expressed genes in the indicated clusters at P25. (E) Violin plots for Cxcl12, Pdgfra, Enpp2, Bmp2, and Timp3 expression in the coronal papilla (cluster 1) cells at P3.5, P7.5, and P25.

### Mx1 and Cxcl12 combination selectively labels pulp progenitor cells

The *in vivo* function of endogenous DPSCs remains elusive due to a lack of mouse models for DPSC fate mapping. In the bone marrow, CXCL12 is highly expressed in reticular cells (CAR cells) and skeletal stem/progenitor cells^29^. Therefore, we next tested whether endogenous DPSCs have a selective expression of CXCL12 in post-erupted molars. By using the *Cxcl12-* GFP mouse model, we first examined which pulp cells express CXCL12 before and after eruption and found that a subset of central pulp cells expressed *Cxcl12-*GFP while mature odontoblasts were *Cxcl12-*GFP negative at P3.5. Interestingly, more pulp cells expressed *Cxcl12-*GFP at P7.5 and the majority of pulp cells highly expressed *Cxcl12*-GFP at P25, indicating that *Cxcl12-*GFP is a reliable marker for adult pulp cells, but not specific to DPSCs (Figure 2A). We previously found that *Mx1-*Cre can inducibly label endogenous skeletal stem cells with high CXCL12 expression in both the periosteum and bone marrow^30,31^. Therefore, to test whether the combination of *Mx1-*Cre and *Cxcl12-*GFP can label endogenous DPSCs, we developed trigenic Mx1/Tom/Cxcl12-GFP reporter mice by crossing *Mx1-*Cre^+^;*Rosa26*-Tomato^f/f^ with *Cxcl12-*GFP^+^ mice. In these mice, treatment with polyinosinic:polycytidylic acid (pIpC: 25 mg/kg) can induce the tdTomato labeling of *Mx1*^+^ cells, which results in yellow *Mx1*^+^*Cxcl12-* GFP^+^ progenitor cells (Tom^+^GFP^+^) and red only differentiated cells. When mice were pIpC-treated at postnatal day 3 and 4 (P3 and P4) and the first molar was analyzed at P7.5, a small subset of *Cxcl12-*GFP^+^ cells (∼2-3%) were *Mx1*-labeled (hereafter *Mx1*^+^) and these double-positive cells exclusively resided in central pulp. Furthermore, the *Mx1*^+^*Cxcl12-*GFP^+^ cell population increased at P25 (∼9-10%) while *Mx1* single-positive cells were enriched in the odontoblast layer (Figure 2B). We next tested the location of PDGFR^+^ pulp progenitor cells by using *Pdgfra-*H2B-GFP mice and found that most pre-odontoblasts are positive for *Pdgfra-* H2B-GFP (Figure 2C). To test whether *Mx1-*Cre labels pre-odontoblasts, we crossed *Mx1-* Cre^+^;*Rosa26*-Tomato^f/f^ mice with *Pdgfra-*H2B-GFP^+^ line. When Mx1/Tom/Pdgfra*-*H2B-GFP mice were pIpC-induced at P3 and P4, there was no detectable *Mx1*^+^ cell contribution to *Pdgfra-*H2B-GFP^+^ in the pulp or odontoblast layer at P7.5, but the *Mx1*^+^*Pdgfra-*H2B-GFP^+^ cell numbers were increased at P25, indicating that early *Mx1*^+^ pulp cells at least minimally contribute to *Pdgfra-* H2B-GFP^+^ pre-odontoblasts (Figure 2D). Interestingly, *Mx1*^+^ cells in dental follicles mainly contributed to periodontal ligament (PDL) cells and without *Cxcl12-*GFP or *Pdgfra*-H2B-GFP expression, suggesting that the *Mx1* and *Cxcl12-*GFP combination can selectively label pulp progenitor cells while *Mx1*^+^ PDL progenitor cells are negative for *Cxcl12-*GFP (Figures S2A and S2B).

**Figure 2.**
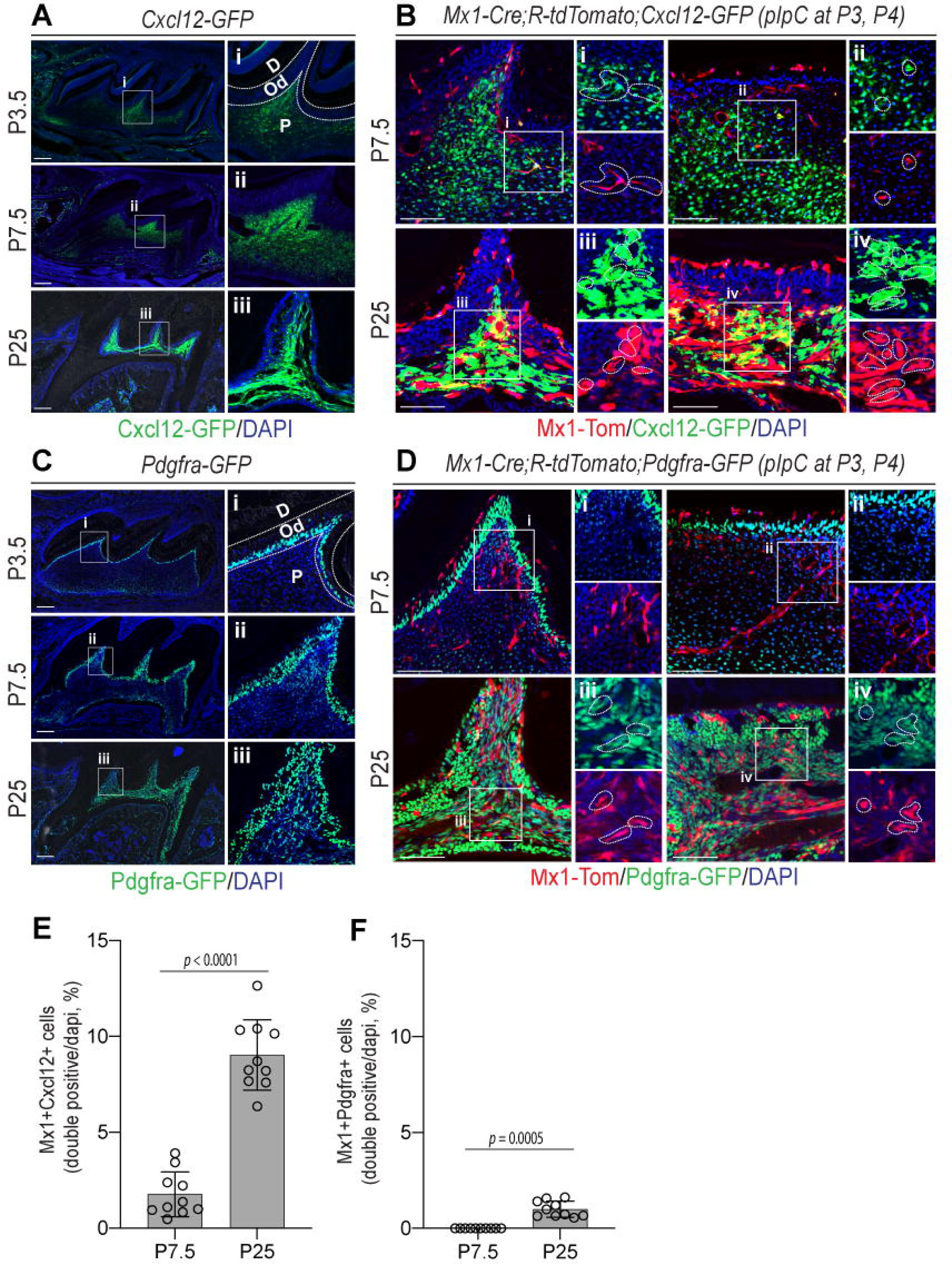
*Mx1* labels a subset of dental pulp cells expressing *Cxcl12*-GFP. (A) Representative fluorescence images of maxillary first molars of *Cxcl12*-GFP reporter mice at P3.5, P7.5, and P25. Dentin (D), Odontoblasts (Od), and Pulp cells (P) in the white boxes (i-iii) are enlarged on the right images. (B) *Mx1*-Cre;*Rosa*-tdTomato;*Cxcl12*-GFP reporter mice were pIpC-induced (P3, P4). Representative fluorescence images of their maxillary first molars at P7.5 and P25 show the pulp horn (left) and pulp chamber roof (right) of molars. The white boxes to the right (i-iv) show enlarged images of *Mx1*^+^*Cxcl12*-GFP^+^ cells (i-iv, yellow dotted lines) with GFP (top) and Tomato (bottom) separation. (C) Representative fluorescence images of maxillary first molars of *Pdgfra*-H2B-GFP reporter mice at P3.5, P7.5, and P25. The white boxes (i-iii) were enlarged on the right side. (D) *Mx1*-Cre;*Rosa*-tdTomato;*Pdgfra*-H2B-GFP reporter mice were pIpC-induced (P3, P4). Representative fluorescence images of their maxillary first molars at P7.5 and P25 show the pulp horn (left) and pulp chamber roof (right) of molars. Enlarged *Mx1*^+^*Pdgfra*-GFP^+^ cells (i-iv, dot lines) in white boxes (i-iv) are shown on the right side with GFP (top) and Tomato (bottom) separation. (n=10). (E, F) Graphs show the double-positive cells for each mouse line (*Mx1*^+^*Cxcl12*^+^ and *Mx1*^+^*Pdgfra*^+^) counted on the pulp and odontoblast layer. Results are displayed as mean ± SEM. Scale bar: 200 μm (A,C), 100 μm (B,D).

### Mx1^+^ pulp progenitor cells are long-term repopulating cells and a main source of adult pulp cells

An important function of DPSCs is their long-term maintenance *in vivo*. To test whether *Mx1*^+^ pulp progenitor cells in mature molars are long-term repopulating cells *in vivo*, Mx1/Tom mice were pIpC-induced at P11 and P13 (post-eruption) and *Mx1*^+^ cells were analyzed at 14 days, 4 weeks, and 16 weeks of age. Three days after pIpC labeling (P14), only a small portion of the pulp cells (∼5% ± 2) in the maxillary first molar were *Mx1*-labeled (Tomato^+^) (Figure 3A, 2 wks, Tom^+^). However, the number and percentage of *Mx1*^+^ pulp cells continuously increased at 8 and 20 weeks of age (∼20% at 8 weeks and ∼80% at 20 weeks of age, respectively) (Figure 3A, 8 wks and 20 wks, Tom^+^). Consistently, we found that over 40% of the pulp cells are *Mx1*^+^ cells in both apical and distal incisors (∼ 40% at 20 wks of age), supporting that *Mx1*^+^ progenitor cells in the early postnatal tooth are the major source of newly generated pulp cells. Notably, we observed the development of a dentinal tubule-like extension from *Mx1*^+^ cells located in occlusal-side odontoblast layers at 20 weeks of age (Figures 3Av and 3Avi, white arrows), which were barely observed at 4 to 8 weeks of age. In addition, the similar dentinal tubules from *Mx1*^+^ cells were observed in the frontal-side (white arrow), but not in the rear-side (root-side) of mandibular incisors (not shown on the Figures), at 20 weeks of age, indicating that the new odontoblast differentiation by the *Mx1*^+^ pulp cells appear on the incisor frontal-side (Figure 3B). By contrast, *Mx1*^+^ cells in the PDL contributed to the majority of PDL cells and alveolar osteoblasts, suggesting that *Mx1*^+^ follicle cells are similar to the previously identified PTHrP^+^ DF progenitor cells (Figures S3A and S3B)^15^.

**Figure 3.**
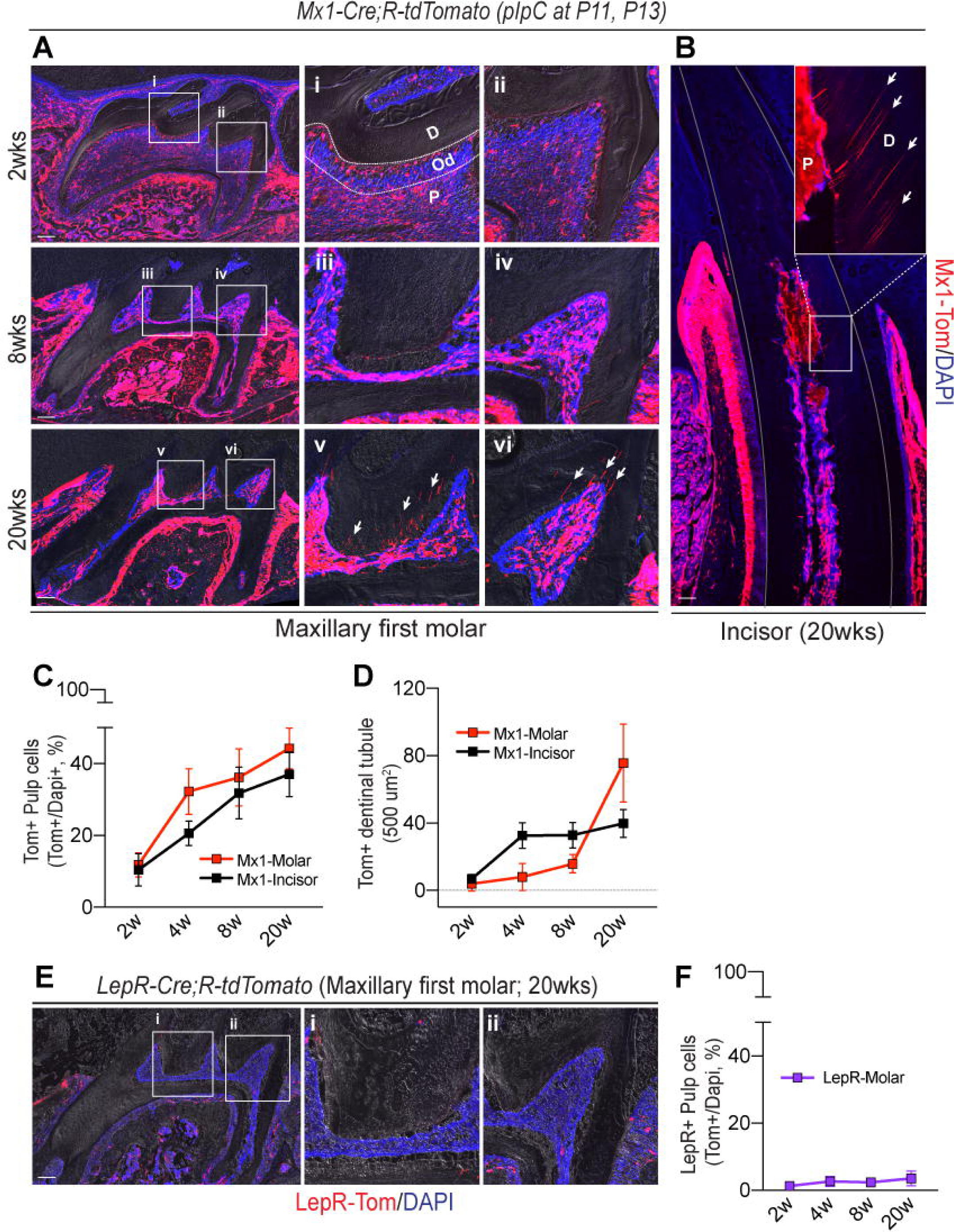
*Mx1*, but not *LepR*, marks postnatal pulp progenitor cells. (A,B) Mx1-Cre;tdTomato reporter mice were pIpC-induced (at P11, P13), and (A) Tomato^+^ pulp cells in maxillary first molars were analyzed at the indicated time points (2 wks, 8 wks and 20 wks). The white boxes (i-vi) indicate enlarged images on the right side. *Mx1*^+^ dentinal tubules (Tom^+^) on dentin area is indicated by white arrows (v,vi). (B) The expression patterns of *Mx1*^+^ cells in the mandibular incisor were determined at 20 weeks of age. The white box is enlarged on the top right, and *Mx1*^+^ dentinal tubules (Tom^+^) are indicated by white arrows. (C) The proportion of *Mx1*^+^ pulp cells in molars and incisors were calculated based on DAPI staining of whole pulpal cells from 2 weeks through 20 weeks. (D) The numbers of Tom^+^ cells in the odontoblast layer (Tom^+^ odontoblast cells) were counted by 500 µm^2^ of each molar and incisor sections (n=5). (E) At 20 weeks of age, *LepR*-Cre;*Rosa*-Tomato reporter mice were analyzed to locate *LepR*^+^ cells in the dental pulp and dentin matrix. White boxes are enlarged on the right side (i,ii) (F) Percentages of *LepR*^+^ cells over DAPI staining within the dental pulp were calculated (n=5). Dentin (D), Odontoblasts (Od) and Pulp cells (P). Results are displayed as mean ± SEM. Scale bar: 200 μm.

While DPSCs are proposed to be a promising cell source for cranial and jaw bone defects, whether endogenous DPSCs have skeletal progenitor characteristics is unknown. We previously found that *Mx1*^+^ SSCs highly overlapped with *LepR*-Cre^+^ SSCs with *Cxcl12-*GFP expression^29^. Therefore, we next tested whether pulp progenitor cells express LEPR by developing *LepR*-Cre^+^;*Rosa26*-Tomato^f/w^ mice. Interestingly, the first and second molars had little or no *LepR*-Cre^+^ cells (Tomato^+^) in the dental pulp and follicles at 2 weeks of age. Even after 20 weeks of age, we observed nearly no detectable *LepR*-Cre^+^ cells in the entire molar pulp, which indicates that *Mx1*^+^ DPSCs are distinct from bone progenitors, with a unique microenvironment regulation and differentiation (Figures 3E and 3F).

### Mx1 selectively labels pulp progenitor cells that contribute to new Ocn^+^ odontoblasts under homeostatic conditions

We next tested whether *Mx1*^+^ pulp progenitor cells contribute to new odontoblasts or whether *Mx1-Cre* can label mature odontoblasts that later develop dentinal tubules. Given that mature odontoblasts were reported to express osteocalcin (*Ocn*)^32^, we developed a trigenic *Mx1*-Cre^+^;*Rosa26*-Tomato^f/w^;*Osteocalcin-*GFP^+^ (Mx1/Tomato/Ocn-GFP) mouse line. In this model, *Mx1*^+^ pulp progenitor cells are Tomato single positive (Tom^+^GFP^-^) and when *Mx1*^+^ progenitor cells differentiate into *Mx1*^+^*Ocn*^+^ odontoblasts, they express Tomato and GFP^33^. Immunofluorescence images of the maxillary first molar (pre-eruption stage) of 2 week-old Mx1/Tomato/Ocn-GFP mice (3 days post-pIpC injection at P11 and P13) showed that the majority of *Mx1*^+^ cells are Tomato single positive and reside in the central pulp. There is no labeling of existing odontoblasts (green), although a subset of *Mx1*^+^ cells is located in the pre-dentin area, which is only observed in premature dentin. (Figure 4A, 2wks). However, at 8 weeks of age, we observed that *Mx1*^+^ cells repopulated in the dental pulp and *Mx1*^+^*Ocn*^+^ double-positive cells appeared in odontoblast cell layers (Figure 4A, 8wks). Notably, at 20 weeks of age, we found a marked increase in the number of *Mx1*^+^ pulp cells (Figure 4D) and *Mx1*^+^ odontoblasts with *osteocalcin*-GFP expression and distinct dentinal tubules (Figure 4A, 20wks, white arrow), suggesting that *Mx1*^+^ pulp progenitor cells move to the odontoblast layer and differentiate into new *Ocn*^+^ odontoblasts. High magnification images (100X) further defined immature *Mx1*^+^ cells within the odontoblast layer at 2 weeks of age (Figure 4B, 2wks, Tom+, arrows) and their *Ocn*^+^ odontoblast differentiation at 20 weeks of age (Figure 4B, 20wks, yellow wedge). Interestingly, these new *Mx1*^+^ odontoblasts are enriched in the loaded plane of the molar (Figures 4B and 4E, occlusal side), but nearly undetectable in the unloaded plane (Figures 4C and 4E, non-occlusal side), suggesting that exogenous mechanical stimuli are required for *Mx1*^+^ DPSC activation and their odontoblast differentiation over time in adult mice. In addition, *Mx1*^+^ cells within the PDL only contributed to *Ocn*^+^ cells in the alveolar bone surface, but not to *Ocn*^+^ cells in cementum layers of PDL at 20 weeks of age (Figures S4A and S4B), further supporting that *Mx1*^+^ DPSCs differ from endogenous PDL progenitors that can differentiate into cementoblasts^15^.

**Figure 4.**
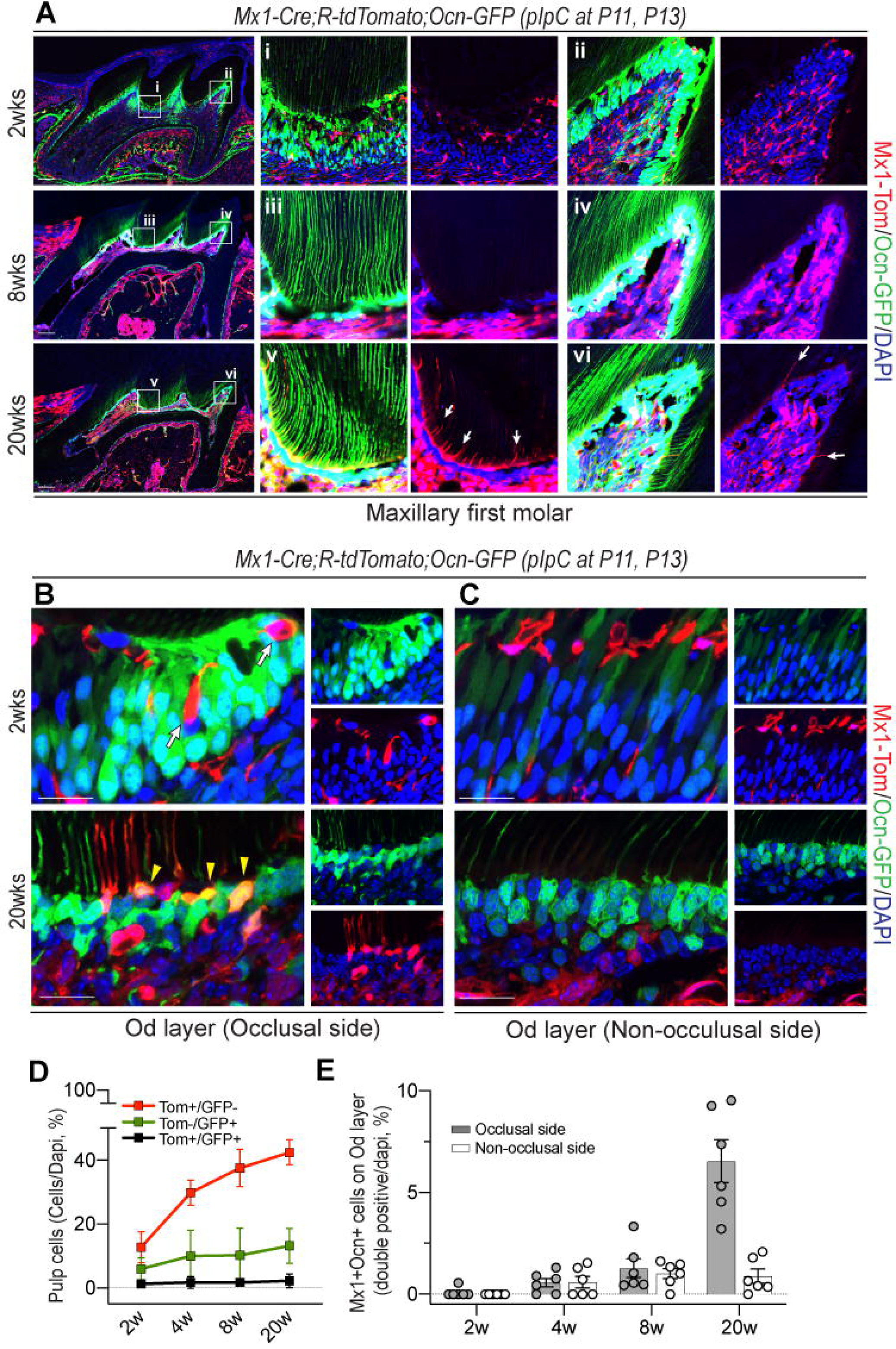
*Mx1*^+^ pulp progenitor cells differentiate into mature odontoblasts. (A,B,C) pIpC-induced (at P11, P13) *Mx1*-Cre;*Rosa*-tdTomato;*Ocn*-GFP reporter mice were analyzed with a fluorescence microscope. (A) Representative images of maxillary first molar at 2, 8, 20 weeks of age with DAPI staining. White boxes were enlarged on the right side (i-vi) and split into Tom-DAPI (right) channels. White arrows show endogenous expression of *Mx1*^+^ cells (v,vi). (B,C) 100X microscope images focused on odontoblast layers at 2 and 20 weeks of age. Each of the images were split into GFP-DAPI (top) and Tom-DAPI (bottom) channels on the right side. (B) Pulp chamber roof (occlusal side) of odontoblast layers include *Mx1*^+^ originating odontoblast cells (black arrow, 2wks) and their mature form (yellow wedge, 20wks) with dentinal process from cell bodies. (C) Odontoblast layers for lateral side of maxillary first molars (non-occlusal side) have less Mx1^+^ originating cell intervention, as well as *Ocn*-GFP expression, on residual odontoblasts compared to the occlusal side. (D) Percentages of Tom^+^/GFP^−^, Tom^−^/GFP^+^, and Tom^+^/GFP^+^ cells over DAPI staining within the dental pulp at different ages were calculated based on sections. (E) The numbers of Tom^+^/GFP^−^, Tom^−^/GFP^+^, and Tom^+^/GFP^+^ cells within the odontoblast layer at different ages were counted based on sections (500 μm^2^, n=5). Results are displayed as mean ± SEM. Scale bar: 200 μm (A,F), 10 μm (B,C), respectively.

### Mx1^+^Ocn*^−^* pulp cells show high expression of dental pulp stem cell markers

DPSCs have been reported to possess many *in vitro* characteristics of mesenchymal stem cells (MSCs) with high clonogenicity and MSC marker expression^34–40^. However, the *in vivo* stem cell properties and physiological function of DPSCs in adult teeth remain elusive. Therefore, we next examined whether *Mx1*^+^ pulp progenitor cells express stem cell markers and their contribution to mature odontoblasts quantitatively. Fluorescence-activated cell sorting (FACS) analysis of molar pulp cells from 2-week old and 20-week old Mx1/Tomato/Ocn-GFP reporter mice revealed that only 5.54% (5.2 ± 0.4, n = 5) of CD45^−^CD31^−^Ter119^−^Lpulp cells were *Mx1* positive at 2 weeks of age. However, this percentage increased almost 4-fold (∼17.4%) at 20 weeks of age (Figure 5A, red box). Furthermore, the percentage of the *Mx1*^+^*Ocn*-GFP^+^ population was increased by ∼2-fold from 2 weeks to 20 weeks of age (Figure 5A, black box), which is consistent with our previous finding of increasing double-positive cells with age (Figure 4A). Subsequent DSPC marker analysis revealed that most (∼80-90%) *Mx1*^+^*Ocn*-GFP^−^ cells expressed DSPC immunophenotypic markers (CD29, CD73, CD90.2, CD146, CD200, Sca1), and this expression was durable even after 20 weeks of age (Figures 5C and 5D). However, control and double-positive cells did not have such high expression of these markers, indicating that DPSCs are enriched in *Mx1*^+^*Ocn*-GFP^−^ cells (Figures 5C and 5D).

**Figure 5.**
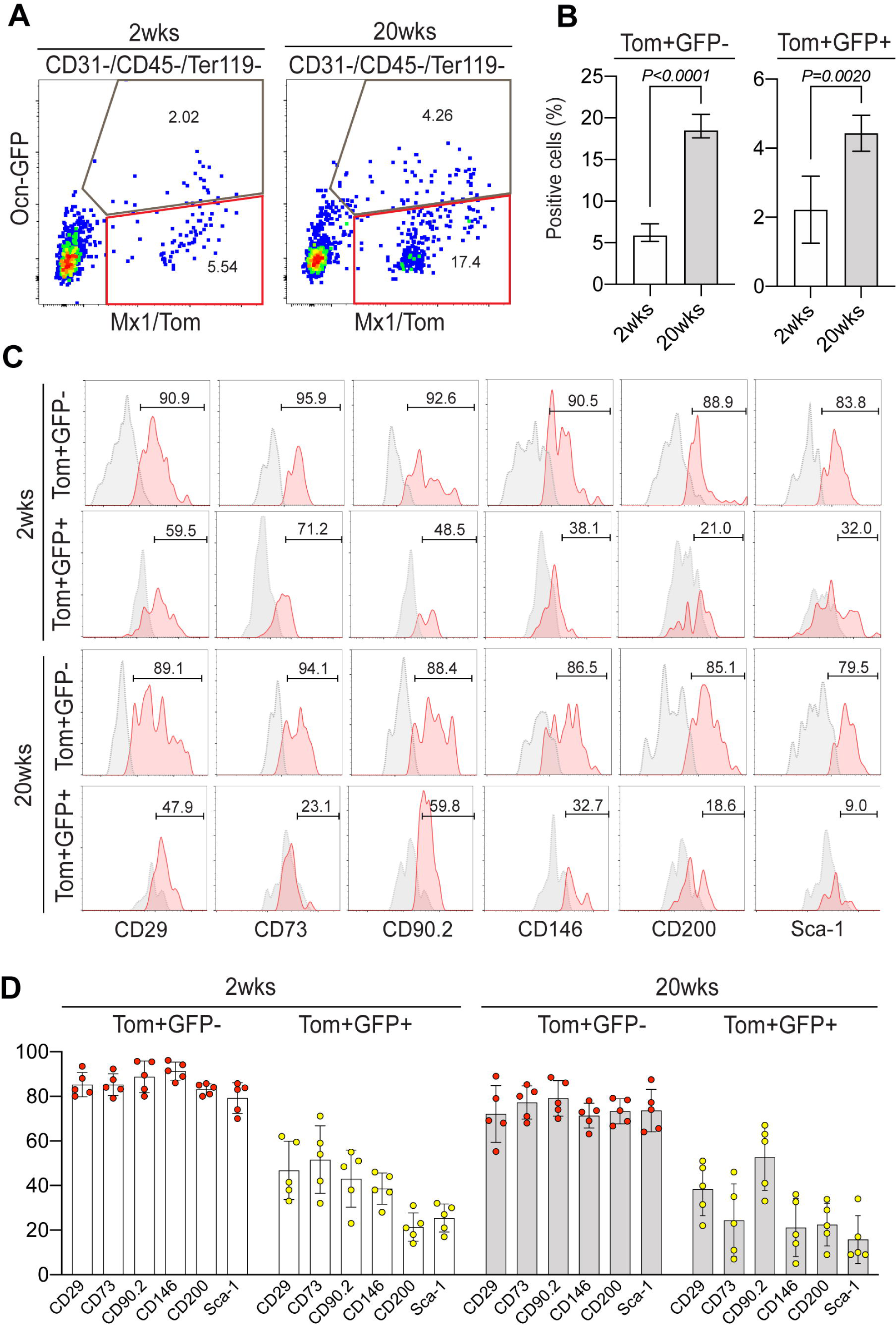
*Mx1*^+^ pulp progenitor cells express DPSCs markers. (A) 2- and 20-weeks old molar pulp cells of *Mx1*-Cre;*Rosa*-tdTomato;*Ocn*-GFP reporter mice were analyzed by FACS with CD31, CD45, Ter119 negative fraction and segregated with Tomato (x-axis) and GFP (y-axis). Red boxes indicate the Tom^+^GFP^−^ population, and grey boxes the Tom^+^GFP^+^ population. Repeatedly analyzed FACS data were collected and represented in (B). Mean ± SEM from three independent experiments with 3–5 mice per session. (C) Relative cell surface expressions of DPSCs markers (CD29, CD73, CD90.2, CD146, CD200, Sca-1) of Tom^+^GFP^−^ cells from 2- and 20-weeks old mice. N=5 for each group.

### Mx1^+^ DPSCs contribute to new odontoblasts in molar dentin injury

Reparative dentin formation is a critical process to protect teeth after injury. However, the origin of new dentin-forming odontoblasts at the injury sites remains unknown. Therefore, to examine whether *Mx1*^+^ pulp progenitor cells contribute to odontoblasts and dentin repair *in vivo*, we generated tooth dentin injuries at the mesial groove on the maxillary first molar of 8-week old Mx1/Tomato/Ocn-GFP mice. We used a hand-operated micro drill bit and generated uniform molar injury without opening the cavity (Figure 6A), which can induce partial odontoblast death at the inner dentin layer of injury points without severe pulp damages. We measured residual dentin thickness (RDT) at the injured site and considered 100-200 μm RDT as a moderate injury and <100 μm as a severe injury. Histological analysis of molars at day 3 post-injury revealed that *Mx1*^+^ cells were increased in the pulp chamber of severe defect sites without odontoblast differentiation (*Ocn-*GFP^−^) (Figure 6B, PS3). Interestingly, at day 7 to day 14 after injury, *Mx1*^+^ progenitor cells started to differentiate into new odontoblasts with developing dentinal tubules in the dentin matrix boundaries at the injury sites (Figure 6B, PS7-PS14). On day 21 post injury, we observed that *Mx1*^+^ odontoblasts became fully polarized with the extension of their dentinal tubules and that a subset of *Mx1*^+^ odontoblasts expressed *Ocn-*GFP (yellow wedges) (Figure 6B, PS21, yellow wedges). This expression was further clarified by high-magnification images (Figure S5A). Furthermore, trichrome staining showed the formation of a reparative dentin-like structure at the injury site with *Mx1*^+^ odontoblasts on day 21 (Figure 6B, PS21, trichrome, dotted line), similar to reparative dentin mineralization in previous studies^41,42^.

**Figure 6.**
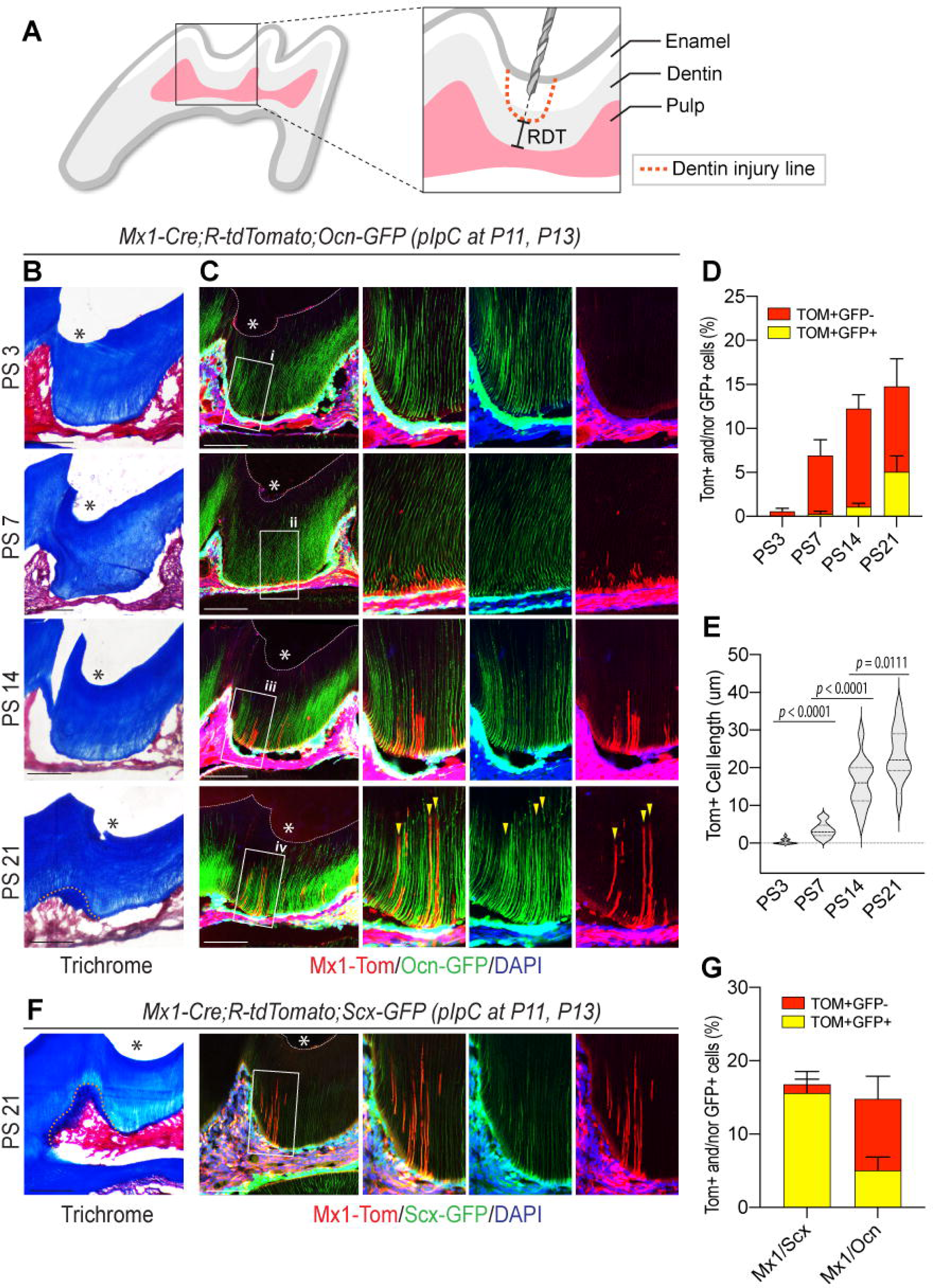
*Mx1*^+^ DPSCs supply the majority of new odontoblasts in dentin injury. (A) Schematic diagram of a site-specific injury point on a maxillary first molar. The dentin injury lines are indicated (orange dashed line) with overall tooth structure. (B,C) A site-specific injury with 200–300 μm RDT depth was formed on the maxillary first molar mesial groove to pIpC-induced (at P11, P13) 4-week old *Mx1*-Cre;*Rosa*-tdTomato;*Ocn*-GFP reporter mice, and responses were analyzed by section imaging from day 3 to day 21 post-surgery (PS 3 to PS 21). (B) Trichrome staining shows the anatomical position of the pulp and dentin as well as the injury point (*) located in the first molar mesial groove for each time point. Yellow dashed lines on PS 21 of trichrome staining data indicate reactionary the dentin line. (C) Fluorescence section images with DAPI staining matching the same slide in B. For each time point, injury points (*) are marked, and the white boxes are enlarged and shown on the right side (i-iv). The enlarged image was split again into different channels with Tom-GFP-DAPI (left), GFP-DAPI (middle), and Tom-DAPI (right). Yellow wedges mark the points where elongated odontoblast process of *Mx1*^+^ originating cells with mature form (iv). (D) The total number of Tom^+^ (GFP^+^ or GFP^−^) cells in the dentinal tubule structure on each 100 μm^2^ section (n=10). (E) The average length of Tom^+^ cells in the dentinal tubules on each 100 μm^2^ section (n=10). (F) A site-specific injury was made on the maxillary first molar mesial groove to pIpC-induced (at P11, P13) 4-week old *Mx1*-Cre;*Rosa*-tdTomato;*Scx*-GFP reporter mice. The *Mx1*^+^ cell response and its overlap with *Scx*-GFP are indicated with trichrome staining (left) and fluorescence imaging (right) on PS 21. Yellow dashed lines on trichrome staining indicate a reactionary dentin line. White box on fluorescence image is enlarged and split into different channels with Tom-GFP-DAPI (left), GFP-DAPI (middle), and Tom-DAPI (right). (G) The comparison expression and cell numbers of Tom^+^GFP^−^ or Tom^+^GFP^+^ cells between *Mx1*^+^ with *Scx*-GFP and *Mx1*^+^ with *Ocn*-GFP mice on PS 21. Each of the counting performed on 100 μm^2^ sections (n=10). Scale bar: 100 μm.

While mature odontoblasts express many osteoblastic genes including Col1, ALP, DMP, and Ocn, they are functionally and morphologically different from bone osteoblasts^43–45^. Interestingly, when we screened for odontoblast gene expression (Figure 1C), we found that mature odontoblasts expressed *screlaxis*, a known tendon/ligament marker. Therefore, we performed an injury experiment using *Mx1*-Cre;*Rosa*-tdTomato;*Scx-*GFP reporter mice and found that *Mx1*^+^ pulp progenitor cells became *Mx1*^+^*Scx-*GFP^+^ odontoblasts with fully polarized dentinal tubules (yellow wedges) with reactionary dentin formation at the site of injury (Figure 6F). Their expression pattern on dentinal tubules was clarified by high-magnification images (Figure S5B). Taken together, these results suggest that *Mx1*^+^ pulp progenitor cells are a main source of fully functional new odontoblasts with polarized dentinal tubules and dentin formation following molar injury.

### Mx1^+^Cxcl12^+^ pulp cells are the most primitive DPSCs and main source for new odontoblasts in adult molars

While *Mx1*^+^ DPSC precursors are the main source of odontoblasts in adult molars, it is possible that *Mx1*^−^ DPSC precursors are present and contribute to new odontoblasts in postnatal molars. Therefore, we performed single-cell sequencing of pulp cells from Mx1/Tomato/Ocn-GFP first molars at P25 (post-eruption stage). UMAP clustering analysis of the single-cell sequencing data showed 6 distinct clusters (Figure 7A). Among these clusters, pulp cell markers and subsequent Tomato (*Mx1*^+^) and GFP (*Ocn*^+^) expression analysis revealed cluster 1 as *Mx1*^+^ CP coronal papilla-like cells, cluster 2 as mid papilla-like cells, cluster 3 as AP (apical papilla)-like cells, cluster 4 as apical follicle cells, cluster 5 as pre-odontoblasts, and cluster 6 as mature odontoblasts, respectively (Figures 7A-7B and S6A-S6B). Notably, Cxcl12 expression highly overlapped with *Mx1* expression and nearly all *Mx1*^+^*Cxcl12*^+^ cells were enriched in cluster 1, suggesting that cluster 1 represents an early DPSC cell population (Figure 7B). This result was supported by monocle pseudotime analysis, which showed that *Mx1*^+^ CP-like cells (cluster 1) are the most with low mineralization gene expression and that they can differentiate to pre-odontoblast, and then odontoblast cells (Figures 7C and S6B). The differentiation trajectory analysis showed that *Mx1*^+^ CP-like cells (cluster 1) have the greatest expression of stem cell markers (i.e. Cxcl12, Pdgfra) with low expression of odontoblast markers (i.e. Osteocalcin, Col1a1) that gradually increased in pre-odontoblast and mature odontoblasts (cluster 6) (Figure 7D). We also observed the presence of distinct *Mx1*^+^*Ocn*^+^ odontoblast clusters, further supporting that *Mx1*^+^ CP-like cells are the main DSPCs that can differentiate into new *Ocn*^+^ odontoblasts in post-erupted permanent molars.

**Figure 7.**
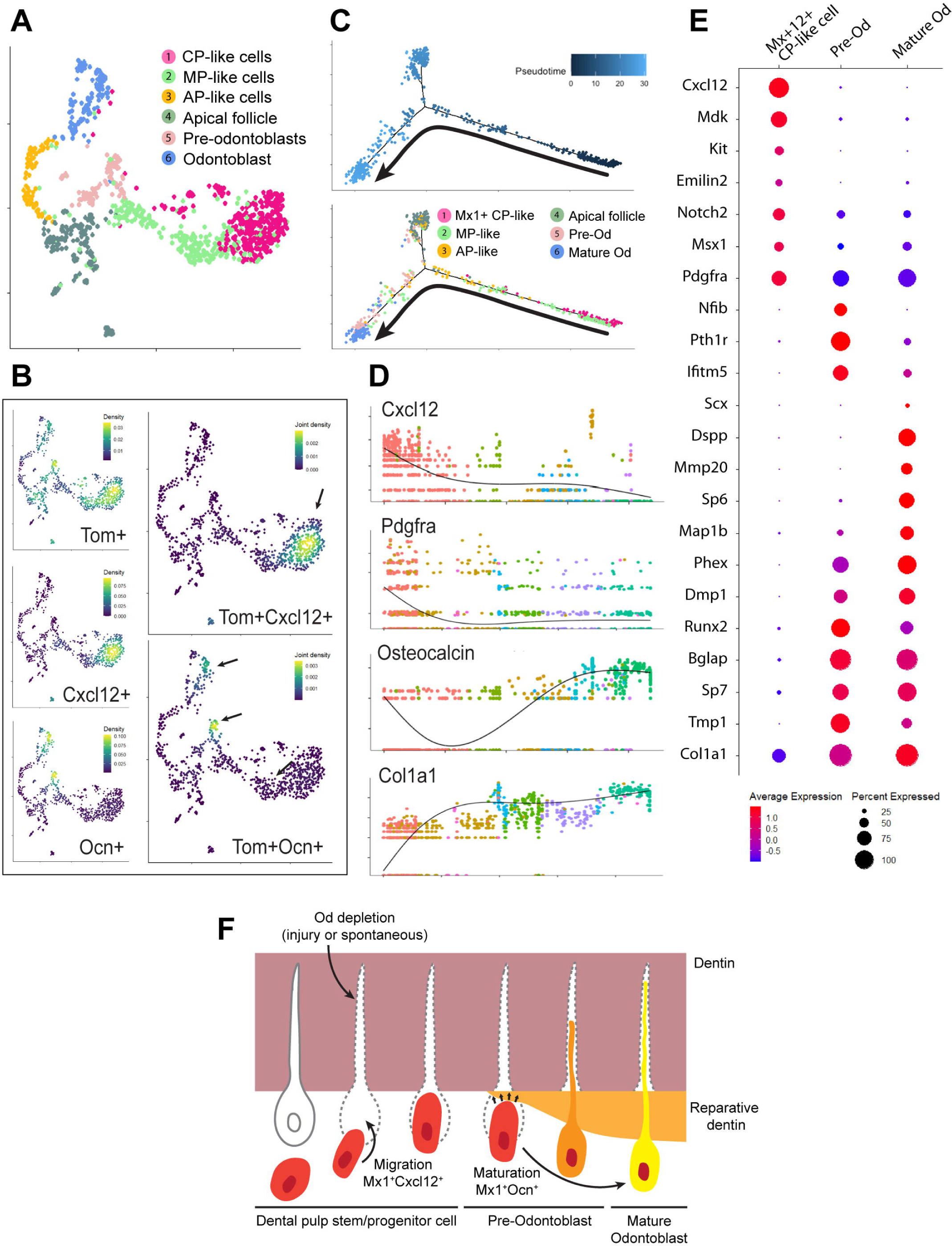
*Mx1*^+^ DPSCs are primitive CP-like cells with the highest expression of stem cell markers. (A) Single-cell analysis of *Mx1*-Cre;*Rosa*-tdTomato;*Ocn*-GFP first molar pulp at P25 and the UMAP visualization of 6 different color-coded clusters of odontoblast related groups. Cluster 1: Coronal papilla, Cluster 2: Middle papilla, Cluster 3: Apical papilla, Cluster 4: Apical follicle, Cluster 5: Pre-odontoblast, Cluster 6: Odontoblast. (B) UMAP-based transcriptional plots for Tomato (Tom^+^), Cxcl12^+^, Ocn^+^ expression and joint plots for Tomato^+^Cxcl12-GFP^+^ and Tomato^+^Ocn^+^ expression. Black arrows on joint plots indicate points of the greatest co-expression. (C) The monocle pseudotime analysis of 6 clusters (top) and the computational model of cell fate decision result with the differentiation trajectory analysis of the different clusters at P25 (bottom). The black arrow is the follow-up of the trajectory of the main cluster according to pseudotime analysis. (D) Monocle pseudotime trajectory expression pattern of representative differentiation genes. X-axis represents pseudotime, and y-axis represents relative expression levels. (E) Gene expression screening for coronal papilla, pre-odontoblast, or mature odontoblast clusters. The analyzed genes were previously identified as stem/progenitor markers for dental pulp (Cxcl12 to Pdgfra) and odontoblast related markers (Nfib to Col1a1). (F) Schematic diagram of migration of *Mx1*^+^*Cxcl12*^+^ cells (dental pulp stem/progenitor cells) and their maturation to *Mx1*^+^*Ocn*^+^ cells (mature odontoblasts).

Further differential gene analysis between *Mx1*^+^ CP-like cells and other pulp cells revealed that *Mx1*^+^ CP-like clusters expressed previously known stem progenitor genes, including Cxcl12, Kit, Notch2, and Msx1. (Figure 7E). Interestingly, we found that *Mx1*^+^ CP-like clusters highly expressed Midkine (Mdk), which can promote odontoblast-like differentiation, although the role of Mdk in adult dental pulp is largely unknown^46^. Taken together, these results suggest that *Mx1*^+^ combined with Cxcl12 can mark endogenous DPSCs, which can differentiate into mature odontoblasts during adult molar homeostasis and injury. These new odontoblasts undergo reparative dentinogenesis and deposit secondary dentin on the bottom of primary dentin (Figure 7E).

## DISCUSSION

Our findings revealed that adult odontoblasts in permanent teeth are continuously replaced by new odontoblasts, and that endogenous pulp progenitor cells maintain the regeneration of pulp cells and new odontoblasts. Our study also demonstrated a unique transition of molar pulp cells, from middle and apical papilla cells in early development to coronal papilla-like cells with *Cxcl12* expression in the post-eruption stage. Notably, a subset of *Cxcl12-*GFP^+^ pulp cells are adult DPSCs that can be labeled by *Mx1*, and these *Mx1*^+^*Cxcl12-*GFP^+^ DPSCs reside in the mid pulp at the early tooth eruption stage, expand, and contribute to the majority of new pulp cells and new odontoblasts forming reparative dentin after injury.

The presence of DPSCs or progenitor cells in adult teeth has long been suggested. Genetic lineage tracing studies with mouse incisor models showed that several distinct mesenchymal populations, including nerve-associated glial cells, periarterial *Gli1*^+^ cells, and *NG2*^+^ pericytes^12,47,48^, contribute to reparative dentinogenesis. Recently, α*SMA*^+^ (α smooth muscle actin) perivascular cells were reported to contribute to reparative dentinogenesis in adult mouse molars^11^. However, many of these studies used tooth injury models; other studies suggested that odontoblasts are post-mitotic cells with certain neuronal features that are normally not replaced during the life of an organism^49–51^. Therefore, it is not clear whether endogenous DPSCs are long-term repopulating cells to supply new odontoblasts or to replace dying odontoblasts under homeostatic conditions. Our study showed that *Mx1* selectively labeled pulp progenitor cells present in the pulp core, but not mature odontoblasts at 2 weeks of age. However, at 20 weeks of age, a marked increase in *Mx1*^+^*Ocn*-GFP^+^ odontoblasts in the dentin matrix in mouse molars was observed (Figure 4). The development and extension of *Mx1*^+^*Ocn*^+^ dentinal tubules later in life further supported that *Mx1*^+^ DPSCs directly differentiate into *Ocn*^+^ odontoblasts. In addition, our data showed that the population of endogenous *Mx1*^+^ pulp progenitor cells remains relatively similar from 4 to 20 weeks, even in the constantly growing incisors, indicating that *Mx1*^+^ pulp progenitor cells include long-term repopulating cells. Our data also suggest that a mild but continuous attrition during mastication or aging (20-week old mouse) is one of the main causes of mature odontoblast depletion; these cells are continuously replaced by new odontoblasts differentiated from *Mx1*^+^ DPSCs rather than through odontoblast proliferation. Single-cell analysis of *Mx1*^+^ pulp cells further supports this finding. While *Mx1*^−^ AP-like and MP-like pulp cells may contribute to some odontoblast differentiation, it is evident that *Mx1*^+^ DPSCs are the most primitive cells in adult molars, and their contribution to mature odontogenesis is increasingly higher than that of other cell clusters. High expression of Mdk in the *Mx1*^+^ CP-like cells suggests a role in adult molar odontogenesis and a potential target for improvement of new odontoblast differentiation (Figure 7D). Further studies will be required to define whether *Mx1*^+^ DPSCs are descendent of neural crest origin mesenchymal cells or derived from peripheral nerve-associated glia cells *in vivo*^12^.

DPSCs and odontoblasts share many osteogenic characteristics and marker expression such as osterix (Osx), osteocalcin (OCN), osteopontin (OPN), collagen type 1 (Col I), and dentin matrix acid phosphoprotein 1 (DMP1), suggesting that DPSCs can be used as a source of transplantation toward alveolar bone defects^43–45^. However, DPSCs and odontoblasts are highly polarized cells with unique features such as dentinal tubule formation, and how they molecularly and functionally differ from osteoblasts has not been elucidated. Our study revealed for the first time that odontoblasts show unique expression of tenogenic marker *Scx-*GFP, although their GFP signal is weaker than those of tendons and PDLs. We also found that *LepR*-Cre, a well-known bone marrow stromal cell marker, does not label DPSCs in mouse molars, indicating that *Mx1*^+^ DPSCs have unique molecular characteristics and can differentiate into *Scx-*GFP^+^ and *Ocn-*GFP^+^ odontoblasts. It remains to be elucidated whether *Scx-*GFP can mark all *Ocn-*GFP^+^ mature odontoblasts or *Ocn-*GFP^−^ pre-odontoblasts.

Our study also provides further insight into the natural healing mechanism in regenerative and reparative dentin formation and provides evidence of the active role of DPSCs in injury response. We used a first molar injury mouse model, generated with a hand-operated micro drill that can inflict physiological and homogenous injuries without heat generation and pulp exposure^11,52^. Interestingly, when the first molar has severe injuries characterized by a remaining dentin thickness (RDT) of 100–200 μm, the *Mx1*^+^ cells migrate up to the dentinal tubule (red wedges) with *Ocn-*GFP and *Scx-*GFP expression observed by day 28 post-injury (Figures 6B and 6F), suggesting that *Mx1*^+^ DPSCs rapidly respond to outer tooth injuries and contribute to new odontoblast-like cells in the replacement of injured odontoblasts.

Taken together, our study supports the notion that there are both intrinsic and extrinsic cues that regulate the functions of adult pulp stem cell subsets, and these DPSCs might have an essential role in maintaining a reparative new odontoblast population with a distinct regulatory mechanism under stress conditions, which could be utilized to improve natural pulp healing and develop a novel therapeutic approach replacing the material base treatment of modern clinical dentistry.

## ACKNOWLEDGEMENTS

This work was supported by the Bone Disease Program of Texas Award and the NIAMS of the National Institutes of Health, under award numbers R61AR078073, R01AR072018, R01DE031288 to D.P and T32DK060445 to L.O and Y.J. The authors are very grateful to L.D. for the valuable drawing of the 3D-printed mouse surgery bed. We thank M.E. Dickinson and J. Kirk in the BCM Optical Imaging and Vital Microscopy Core.

## STAR*METHODS

### Animals

C57BL/6, *Mx1-*Cre^53^, *Rosa26*-loxP-stop-loxP-tdTomato (Rosa-Tom)^54^, and *LepR-*Cre^55^ mice were purchased from The Jackson Laboratory. *Osteocalcin-*GFP (*Ocn-*GFP, C57/BL6 background) mice were kindly provided by Drs. Ivo Kalajzic and Henry Kronenberg. *Cxcl12-*GFP mice were kindly provided by Dr. Nakasawa. Genotyping of all Cre-transgenic mice and GFP fluorescent conjugated mice was performed by PCR (GenDEPOT) using primers for each sequence. Genotyping of the Rosa and GFP locus was performed according to The Jackson laboratory’s protocols. For *Mx1*-Cre induction, mice were injected twice intraperitoneally with 25 mg/kg of pIpC (Sigma) given at two-day intervals. All mice were maintained in pathogen-free conditions, and all procedures were approved by Baylor College of Medicine’s Institutional Animal Care and Use Committee (IACUC).

### Histological analysis

The maxilla was dissected and fixed in 4% paraformaldehyde overnight at 4°C, followed by decalcification in 10% EDTA for 7 days. Decalcified tissue was placed in a 10% sucrose solution overnight and moved to a 30% sucrose solution for storage. The samples were embedded in O.C.T. Compound (Tissue-Tek, SAKURA) before sectioning. Cryosectioning was performed with a cryostat (CM3050S, Leica Biosystems) with the CryoJane tape transfer system (Leica Biosystems). For sections labeled with fluorescent reporters, the slides were stained with DAPI and visualized under the confocal microscope (A1R-s, Nikon) with filter cubes optimized for tdTomato, green fluorescent protein (GFP), and DAPI variants. Trichrome staining was performed using the trichrome stain kit (ab150686, Abcam), according to the manufacturer’s instructions, and imaged by a bright field microscope (Ci-L, Nikon).

### Mouse tooth injury

Mice were anesthetized with an intraperitoneal injection of an anesthetic combination of ketamine (56.3 mg/kg), xylazine (2.9 mg/kg), and acepromazine (0.6 mg/kg) and prepared for the injury procedure on the maxillary first molars. All procedures for molar injury were performed on a 3D-printed surgery bed^56^. A class I cavity was prepared with hand-operated micro drill bits (diameter, 0.20 mm) on the center of the mesial groove of the maxillary first molars ^57^. According to the depth through which the drill bit penetrates, up to 0.1 mm is defined as a shallow cavity, and 0.1–0.2 mm is defined as a deep cavity. After reaching the appropriate depth, the cavity was washed with PBS to remove debris produced by cavity preparation and capped using light-cured composite resin (Flowable, Pentron) associated with a universal adhesive system (3M).

### Statistical Analysis

All measurements were made by a blinded examiner at 3 independent trials, and the average was recorded. All data were expressed as mean ± SEM. For comparison between 2 independent groups, statistical differences were evaluated by unpaired 2-tailed Student’s t-test. One-way analysis of variance, followed by Tukey’s post hoc test, was performed for multiple comparisons. A P value <0.05 was considered as statistical significance.

### Flow cytometry analysis

To analyze the expression of mouse DPSC markers, the mouse head was dissected, and all maxillary and mandibular molars were extracted (first, second and third molars). After grinding the tooth in a mortar to separate the pulp tissue, the tissues were incubated with 5–10 mL of 0.1% collagenase and 10% FBS in PBS at 37°C for one hour. The dissociated pulp cells were washed with PBS and filtered with a 40-mm strainer. Then the cell suspension was stained with mouse CD45-Pacific blue (clone: S18009F), CD31-Pacific blue (clone: 390), Ter119-BV605 (clone: TER-119), CD90.2-APC/Cy7 (clone: 30-H12), CD73-PE/Cy7 (clone: TY/11.8), CD29-APC (clone: HMb1-1), Sca1-APC/Cy7 (clone: D7), CD200-APC (clone: OX-90), and CD146-PE/Cy7 (clone: ME-9F1). Flow cytometry experiments were performed using the LSRII and FACS Aria cytometer (BD Biosciences, San Jose, CA). Data were analyzed with the FlowJo software (TreeStar, Ashland, OR) and represented as histograms, contour, or dot plots of fluorescence intensity.

### Single-cell RNA sequencing analysis

#### Cell Isolation and sequencing

Molars of P25 mice were digested to obtain the single-cell transcriptomes. Briefly, the molars of 3 mice were digested in 4 mg/mL Dispase and 2 mg/mL Collagenase I on a thermomixer at 37°C for 30 min to 1 hour, depending on the stage of the sample, to release the cells from the tissue. For each sample, 10,000 cells were targeted for scRNA sequencing, with the actual sequenced cells at 11,058 cells. Quality control, mapping, and count table assembly of the library were performed using the CellRanger pipeline version 6.1.2.

#### Variable genes and dimensionality reduction

Raw read counts from the cells at each stage were analyzed using the Seurat 4.3R package. Data from P3.5 and P7.5 molars in the literature^58^ were merged and integrated with the P25 dataset. Cells with low gene expression were filtered out following standard Seurat object generation. Cells with <300 genes per cell and with more than 25% mitochondrial read content were filtered out. For the merged datasets, the PrepSCTIntegration function was performed before identifying anchors with the function FindIntegrationAnchors. Seurat objects were returned by passing these anchors to the IntegrateData function. RunPCA and RunUMAP visualizations were used for downstream analysis and visualization. For analysis of the P25 dataset only, Sctransform was applied for normalization and cell cycle regression. RunPCA and RunUMAP were performed for dimensionality reduction and final visualization of the clustering.

#### Subcluster analysis

Heterogeneity within the dental mesenchymal populations was investigated through subcluster analysis. Published markers for the dental mesenchyme were used to screen and identify the different dental mesenchymal cell populations.

For the integrative analysis of samples at different stages, Seurat 4 was used to combine the single-cell data from five stages and perform integration analysis. The PrepSCTIntegration function was run before identifying anchors with the function FindIntegrationAnchors. Seurat objects were then returned by passing these anchors to the IntegrateData function. RunPCA and RunUMAP visualization were used for downstream analysis and visualization.

#### Monocle trajectory analysis

Monocle was used for pseudotime trajectory inference across the dental mesenchymal cells. Monocle inferred cluster and lineage relationships within a given cell type through the inputted cells. The estimation of the root node of the trajectory was based on the stem cell markers such as Cxcl12 and Pdgfra.

