## Supplementary figures and images for "*Mx1*-labeled pulp progenitor cells are main contributors to postnatal odontoblasts and pulp cells in murine molars"

### Supplemental Figure 1

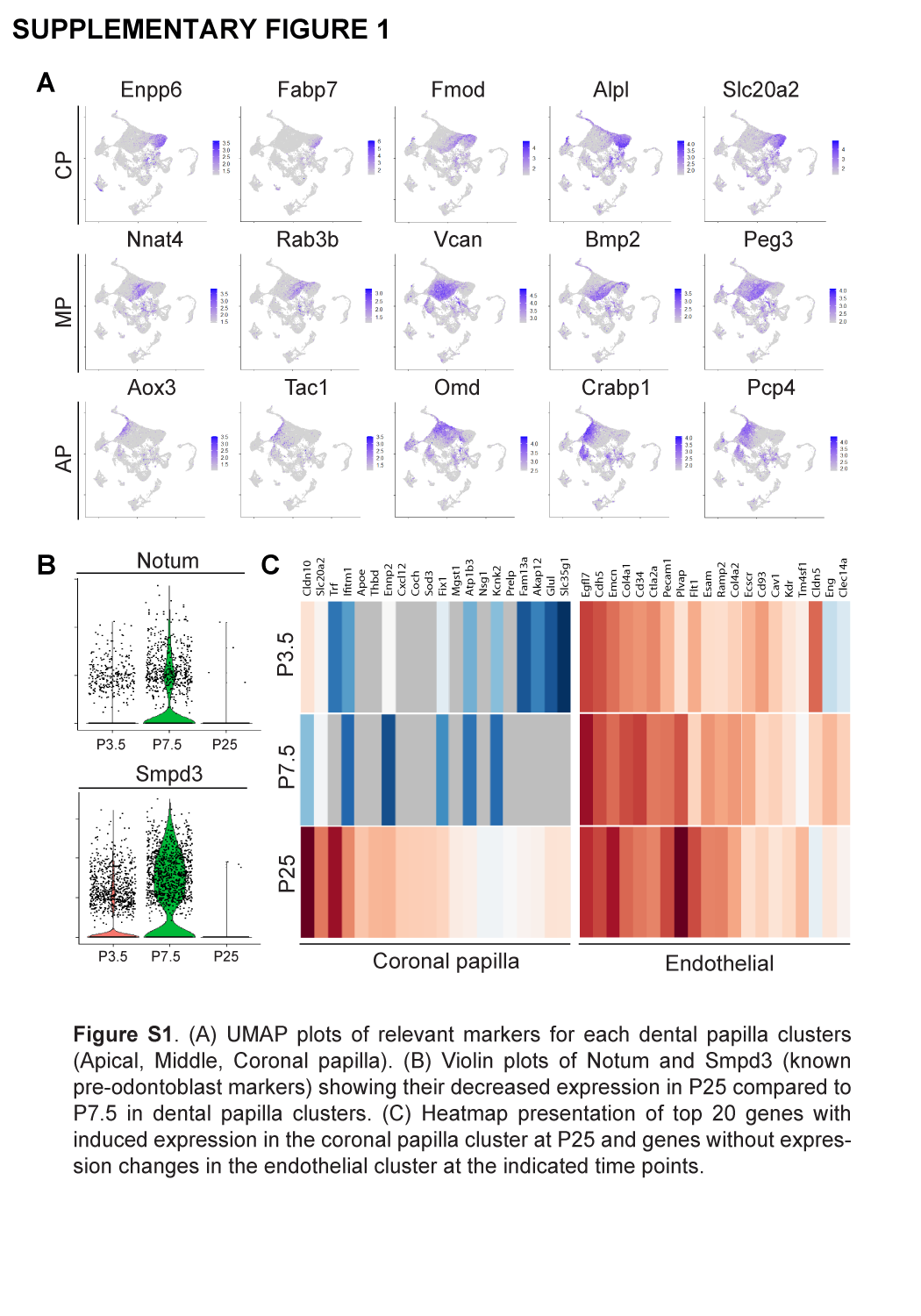

### Supplemental Figure 2

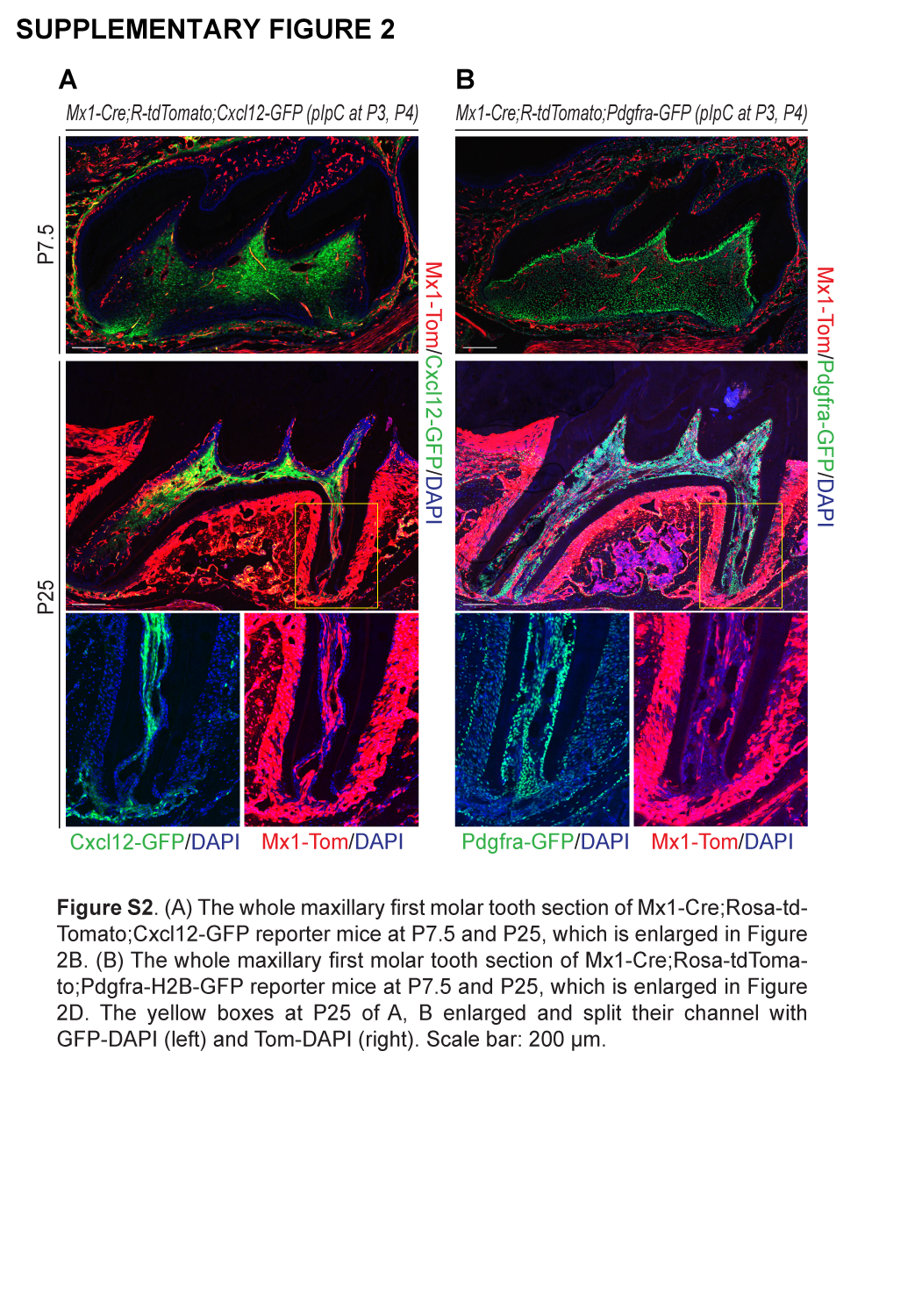

### Supplemental Figure 3

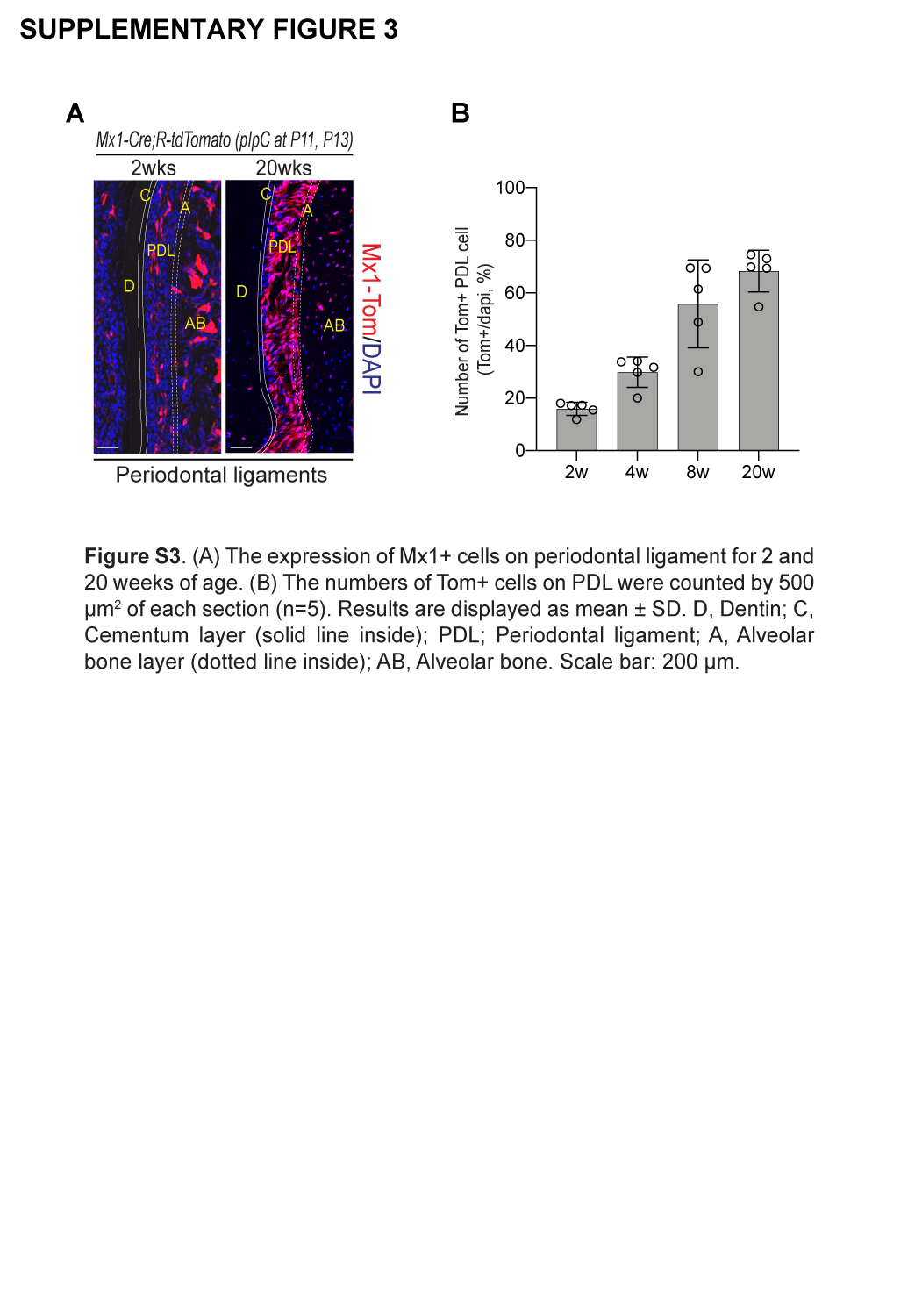

### Supplemental Figure 4

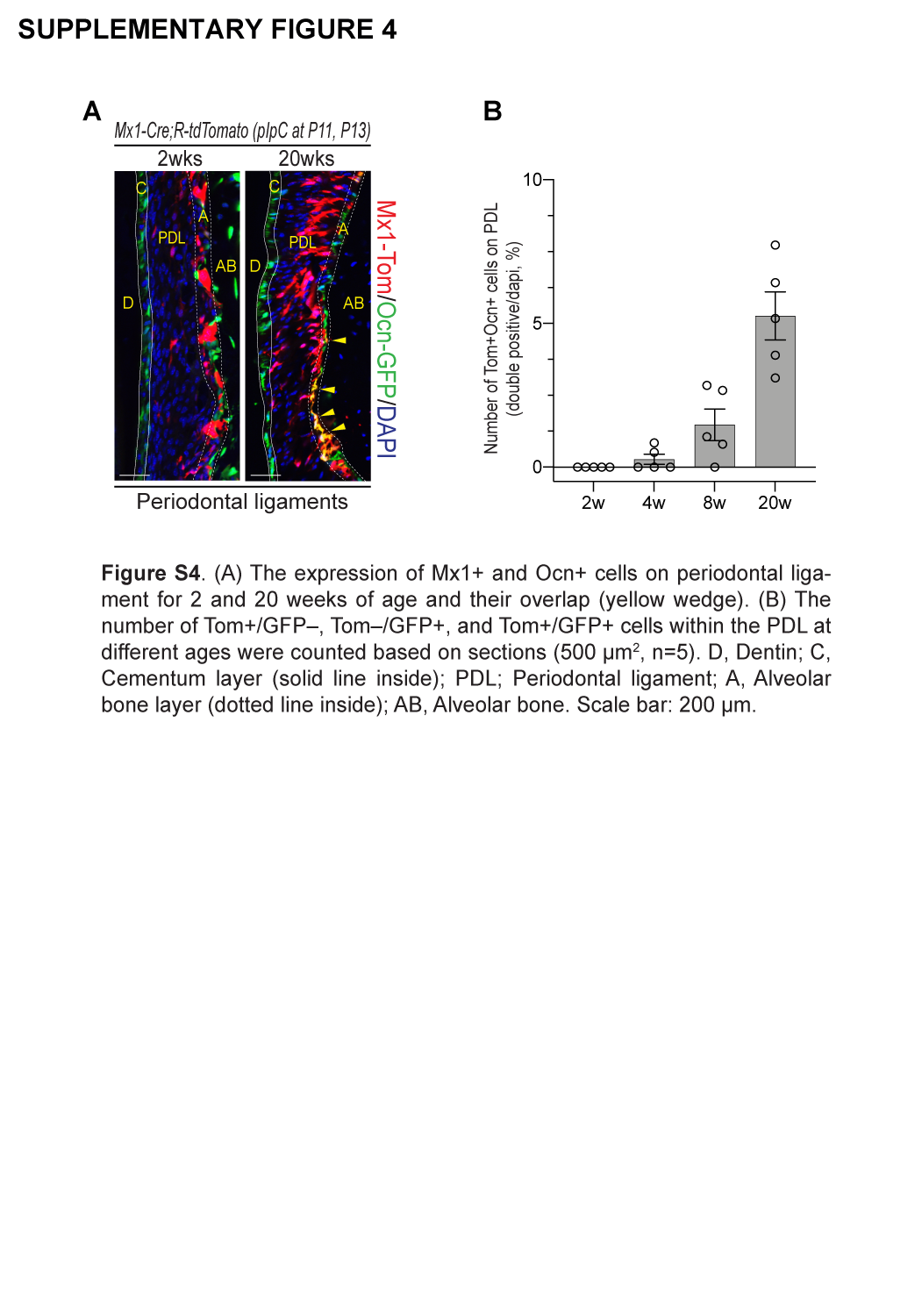

### Supplemental Figure 5

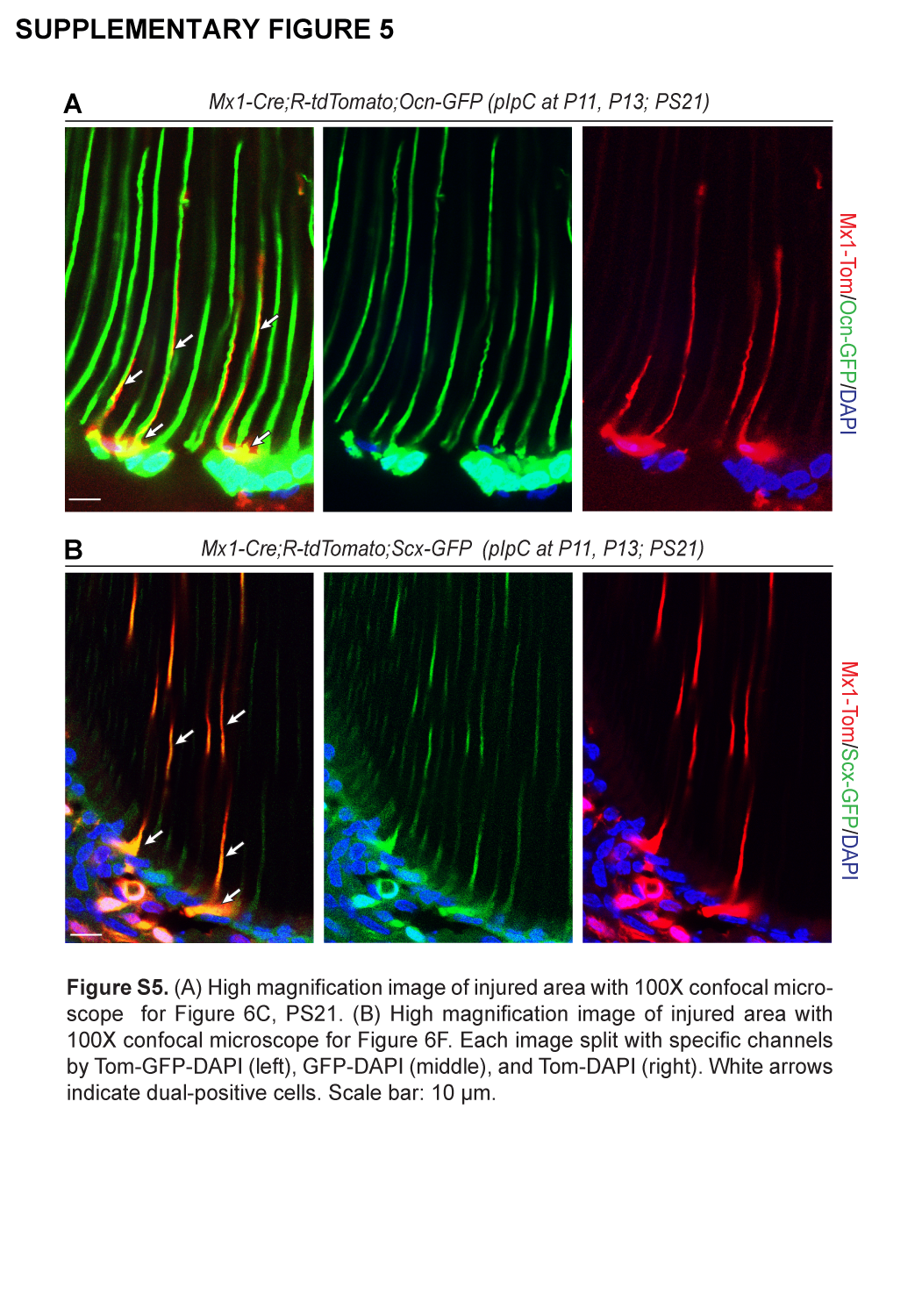

### Supplemental Figure 6

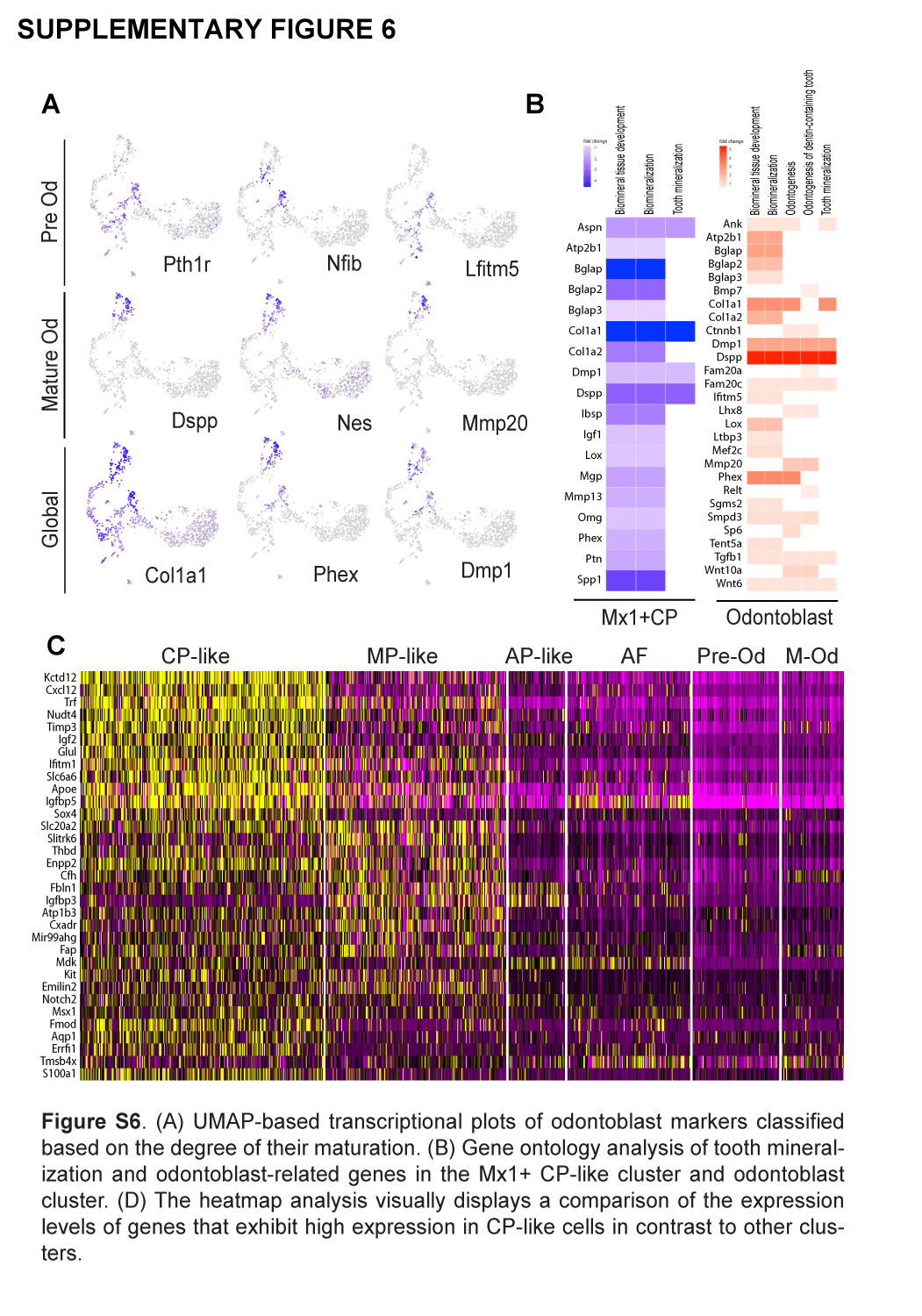
